# Atomic Structures of Glucose, Fructose and Sucrose and Explanation of Anomeric Carbon

**DOI:** 10.1101/002022

**Authors:** Raji Heyrovska

## Abstract

Presented here are the structures of three biologically important sweet sugars, based on the additivity of covalent atomic radii in bond lengths. The observed smaller carbon-oxygen distances involving the ‘anomeric’ carbons of the open chain hexoses are explained here, *for the first time*, as due to the smaller covalent double bond radii of carbon and oxygen than their single bond radii in the cyclic forms and in sucrose. The atomic structures of all the three carbohydrates, drawn to scale in colour, have been presented here also *for the first time*.

## Introduction

Sugars play important roles in our biological functions and well-being [1]. They are digestible for those with normal functions of insulin, but are not for those with diabetes whose insulin function is impaired [2,3]. Now that the World Diabetes Day [4] brings their awareness, the author has made contribution here to the understanding of the structure of these compounds at the ultimate atomic level.

In recent years it was found that bond lengths in simple as well as complex inorganic, organic and biological compounds are exact sums of the appropriate radii of the adjacent atoms and or ions, see e.g., [5–9]. In biological molecules, the known bond lengths in the skeletal structures could be explained as sums of the covalent atomic radii (R_A_), defined as [10] half the bond length, d(AA) between two atoms (A) of the same kind. The linear correlations of known bond lengths with the sums of the appropriate radii were demonstrated for DNA, amino acids, vitamin B2, and many other molecules [5–9]. Here, the atomic structures for the three important sweet carbohydrates, glucose, fructose and sucrose have been presented.

## D-Glucose and D-fructose

The conventional structures [11a,b] of these molecules in open chain forms are shown in Figs. 1a and b. The atoms, carbon (C), oxygen (O) and hydrogen (H) are shown as grey, orange and green circles, respectively, with their respective covalent radii [5–10] in Fig. 2. Note that the double bond covalent radii, C_db_ and O_db_ are smaller than the single bond radii, C_sb_ and O_sb_ (subscripts db = double bond and sb = single bond).

**Fig. 1.**
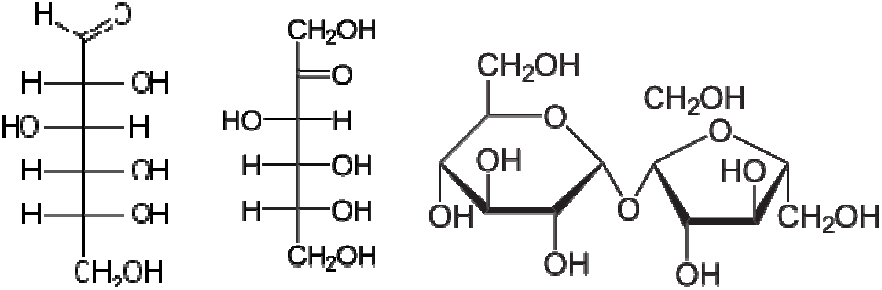
Open chain conventional structures of **a)** D-glucose, [11a] **b)** D-fructose, [11b] and **c)** sucrose, [11c].

**Fig. 2.**
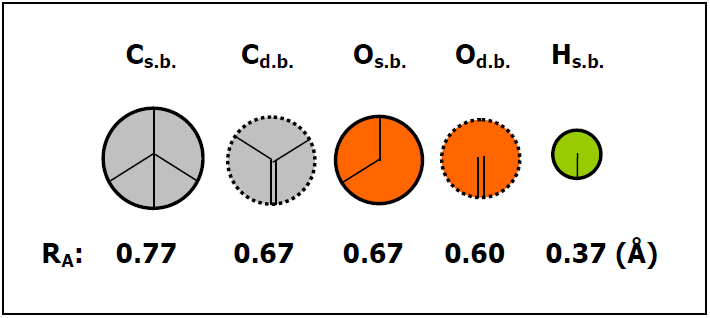
The covalent radii, R_A_ = d(AA)/2 of atoms (A) in Å, [5–10]. Carbon: C (grey), oxygen: O (orange) and hydrogen: H (green). Subscripts s.b. and d.b. stand for single bond (A-A) and double bond (A=A) respectively.

The open chain structures of glucose and fructose, drawn to scale with the covalent radii in Fig. 2, are shown in Figs. 3 a and b. Table 1 gives the bond lengths in these compounds as sums of the appropriate radii of the atoms constituting the bonds. The anomeric [1,12,13] carbons, C(1) of glucose and C(2) of fructose have the double bond radius, C_db_. The oxygen atoms bonded with these carbons also have the double bond radius, O_db_. Hence the corresponding carbon-oxygen distances (1.27 Å) are smaller than the C-O single bond distances (1.44 Å) (see Table 1). This explains for the first time the earlier reports, e.g., 1.24 A and 1.44A [14] for the shorter and longer CO bonds.

**Table 1.**
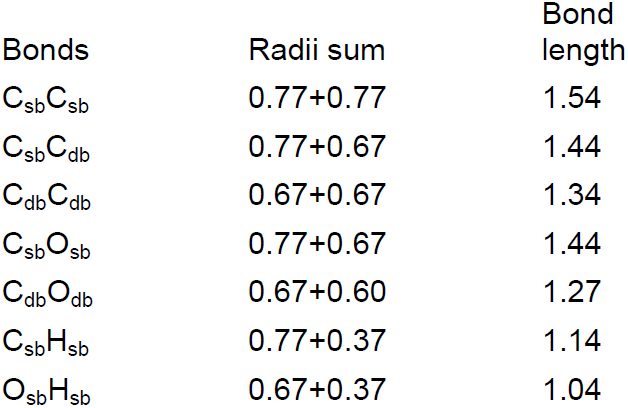
Bond lengths as sums of covalent atomic radii (+/-0.02Å). Subscripts sb: single bond, db: double bond

**Fig. 3.**
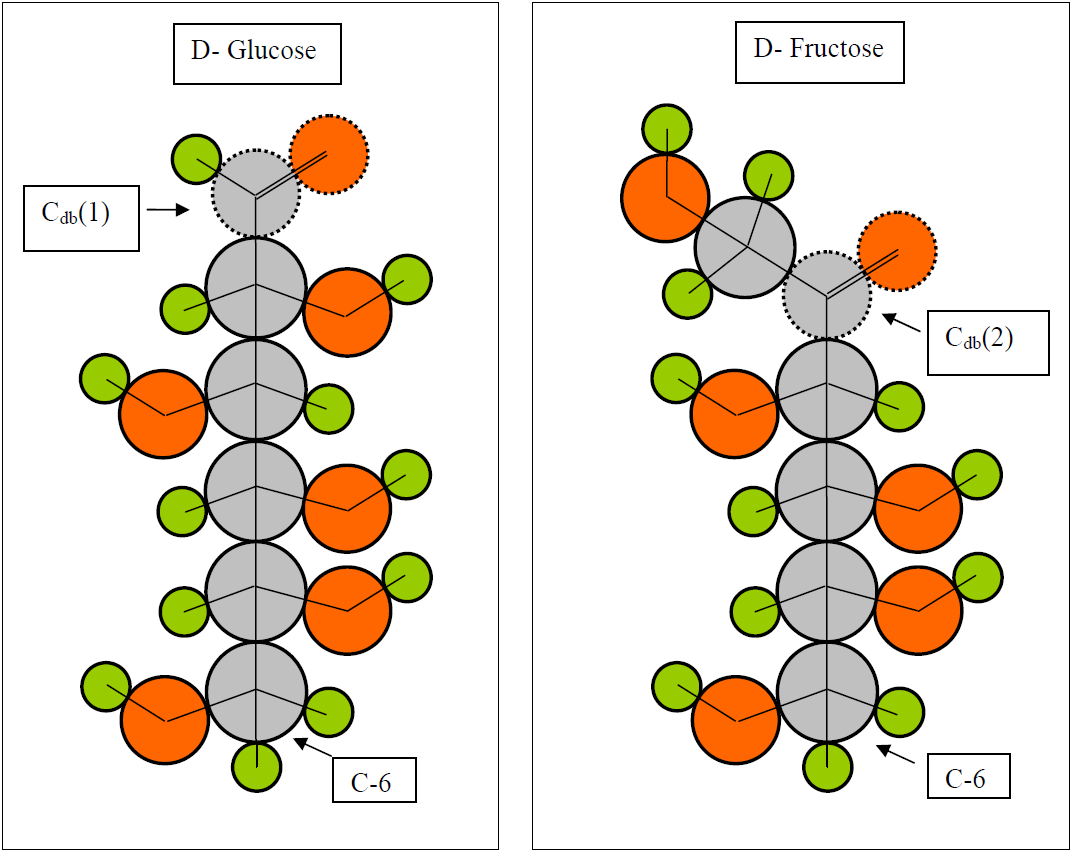
Atomic structures of **a)** D-glucose and **b)** D-fructoseand based on additvity of atomic radii in bondlengths. Note: It is pointed out here for the first time that the ‘anomeric’ carbons C_d.b._(1) of glucose and C_d.b._(2) of fructose have double bond radius, C_d.b._ (= 0.67 Å) and all the other carbons have single bond radius, C_s.b._ (= 0.77 Å). All atoms are drawn to scale as per the radii in Fig. 2 and the various bond lengths are radii sums as given in Table 1.

## Sucrose

Sucrose is conventionally represented e.g., [11c] as in Fig. 1c. The open chain forms with the unsaturated carbons C_db_(1) in glucose and C_db_(2) in fructose become more stable by cyclization with the C(5) carbons and turn into the saturated single bond carbons with larger radii, C_sb_(1) and C_sb_(2) and single bond oxygens with larger radii as well. These are shown in Figs. 4a and b. Glucose forms a pyranose hexagon whereas fructose forms a furanose pentagon.

**Fig. 4.**
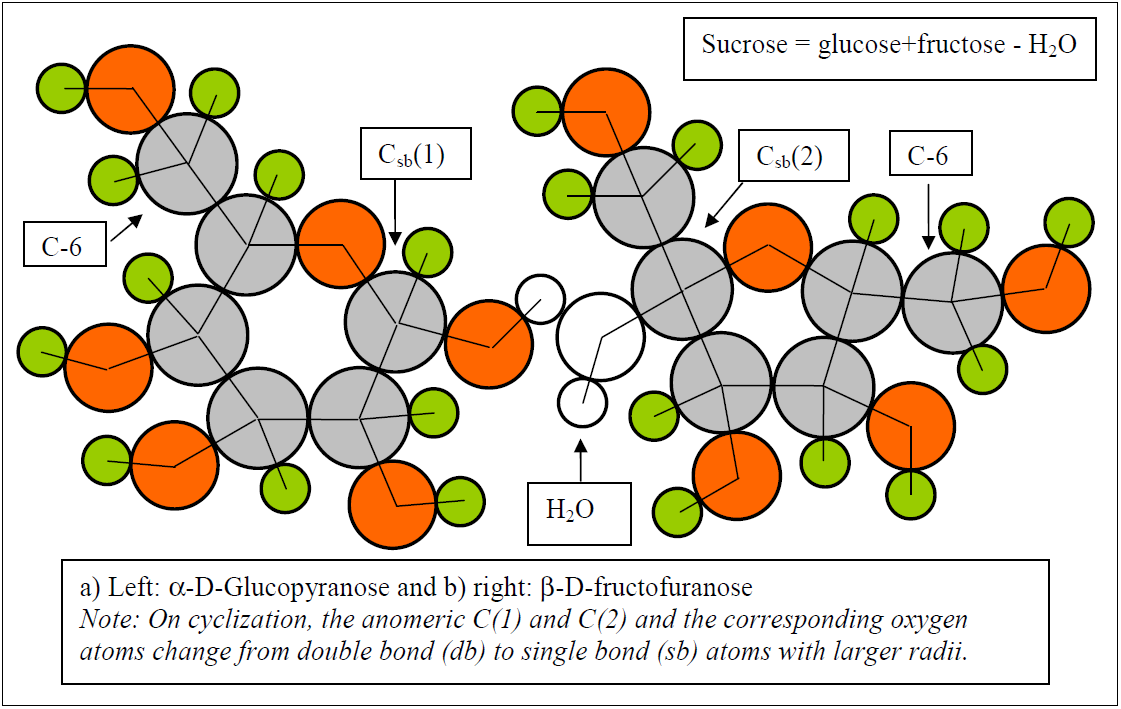
The cyclic forms of D-glucose and D-fructose, namely, D-glucopyranose (**a**: left) and D-fructofuranose (**b**: right). These are more stable since the unsaturated anomeric carbons which have double bond character change into the saturated single bond carbons, C_s.b._(1) and C_s.b._(2), with larger radius. These cyclic forms link together by condensation via formation of a water molecule which readily detaches and form sucrose. All atoms are drawn to scale as per the radii in Fig. 2 and the various bond lengths are radii sums as given in Table 1.

When these two molecules combine by condensation and loss of a water molecule, a glycosidic bond is formed between the anomeric carbons, C_sb_(1) and C_sb_(2) via an oxygen, thereby forming a molecule of sucrose, see Fig. 5. For the spatial locations of the various atoms, see [11d]. Observed [15] carbon-carbon distances are 1.51 - 1.53 Å; carbon-oxygen 1.40 to 1.44 Å; carbon-hydrogen 1.08 to 1.11 Å; oxygen-hydrogen 0.94 to 0.99 Å. These compare well with the bond lengths in Table 1 here. Note that the short anomeric C=O distance found in the open chain forms do not occur in the cyclic forms and in sucrose [15a,b], confirming that all the C-O bond lengths in sucrose involve single bond radii.

**Fig. 5.**
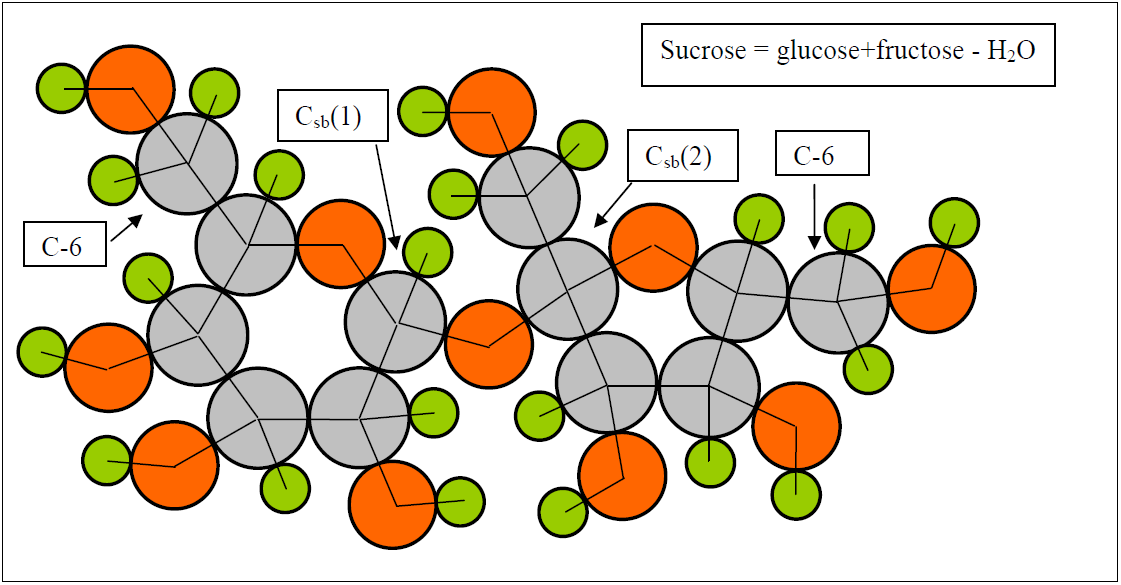
Atomic structure of sucrose. All atoms are drawn to scale as per the radii in Fig. 2 and the various bond lengths are radii sums as given in Table 1.

Thus, this paper presents *for the first time* the precise structures of glucose, fructose and sucrose at the ultimate atomic level and explains the shorter ‘anomeric’ carbon to oxygen bonds of the open chain structures.

